# A clonally reproducing generalist aphid pest colonises diverse host plants by rapid transcriptional plasticity of duplicated gene clusters

**DOI:** 10.1101/063610

**Authors:** Thomas C. Mathers, Yazhou Chen, Gemy Kaithakottil, Fabrice Legeai, Sam T. Mugford, Patrice Baa-Puyoulet, Anthony Bretaudeau, Bernardo Clavijo, Stefano Colella, Olivier Collin, Tamas Dalmay, Thomas Derrien, Honglin Feng, Toni Gabaldón, Anna Jordan, Irene Julca, Graeme J. Kettles, Krissana Kowitwanich, Dominique Lavenier, Paolo Lenzi, Sara Lopez-Gomollon, Damian Loska, Daniel Mapleson, Florian Mauster, Simon Moxon, Daniel R. G. Price, Akiko Sugio, Manuella van Munster, Marilyne Uzest, Darren Waite, Georg Jander, Denis Tagu, Alex C. C. Wilson, Cock van Oosterhout, David Swarbreck, Saskia A. Hogenhout

**Author notes:** These authors contributed equally to the work (Thomas C. Mathers, Yazhou Chen). Current address: GJK: Rothamsted Research, Harpenden, Hertforshire, ALF5 2JQ, UK; KK: J. R. Simplot Company, Boise, Idaho, USA; PL: Alson H. Smith Jr. Agriculture and Extension Center, Virginia Tech, Virginia Tech, Winchester 22602, USA; SLG: Department of Plant Sciences, University of Cambridge, Downing Street, Cambridge CB2 3EA, UK; DRGP: Moredun Research Institute, Pentlands Science Park, Bush Loan, Penicuik, Midlothian, EH26 0PZ, UK; AS: INRA, UMR 1349 IGEPP (Institute of Genetics Environment and Plant Protection), Domaine de la Motte, 35657 Le Rheu Cedex, France.

## Abstract

**Background:** The prevailing paradigm of host-parasite evolution is that arms races lead to increasing specialisation via genetic adaptation. Insect herbivores are no exception, and the majority have evolved to colonise a small number of closely related host species. Remarkably, the green peach aphid, *Myzus persicae*, colonises plant species across 40 families and single *M. persicae* clonal lineages can colonise distantly related plants. This remarkable ability makes *M. persicae* a highly destructive pest of many important crop species.

**Results:** To investigate the exceptional phenotypic plasticity of *M. persicae*, we sequenced the *M. persicae* genome and assessed how one clonal lineage responds to host plant species of different families. We show that genetically identical individuals are able to colonise distantly related host species through the differential regulation of genes belonging to aphid-expanded gene families. Multigene clusters collectively up-regulate in single aphids within two days upon host switch. Furthermore, we demonstrate the functional significance of this rapid transcriptional change using RNA interference (RNAi)-mediated knock-down of genes belonging to the cathepsin B gene family. Knock-down of cathepsin B genes reduced aphid fitness, but only on the host that induced up-regulation of these genes.

**Conclusions:** Previous research has focused on the role of genetic adaptation of parasites to their hosts. Here we show that the generalist aphid pest *M. persicae* is able to colonise diverse host plant species in the absence of genetic specialisation. This is achieved through rapid transcriptional plasticity of genes that have duplicated during aphid evolution.

## Background

Parasites often exhibit a high degree of specialisation to a single or reduced range of host species [1, 2]. This is especially true for insect herbivores, of which there are around 450 thousand described species living on around 300 thousand species of vascular plants, the majority of which are monophagous or oligophagous, being able to colonise only one or a few closely related plant species [3]. Acute specialisation of parasites is likely due to the complex relationships that occur between the parasites and their hosts, with increasing specialisation being driven by coevolutionary arms races [4, 5]. In the case of herbivorous insects, the plant-insect interface represents a dynamic battle ground between host and parasite in which insect effector genes evolve to subvert plant defences and plant resistance genes evolve to detect infection and guide plant immunity [6, 7].

Despite the tendency for parasites to evolve highly specialised relationships with their hosts, occasionally, genuine generalist species with broad host ranges have evolved. For example, clonally produced individuals of the parasitic trematode *Maritrema novaezealandensis* are able to colonise a broad range of crustacean species [8] and the giant round worm *Ascaris lumbricoides,* which causes Ascariasis and infects an estimated 0.8 billion people worldwide , is able to infect both humans and pigs [9]. Often, however, generalist parasite species have turned out to be cryptic specialists, made up of host adapted biotypes or cryptic species complexes [10-12]. For example, the pea aphid *Acyrthosiphon pisum* is considered polyphagous, being found on most plants of the Fabaceae, but actually consists of different biotypes on a continuum of differentiation that colonise specific species of this plant family [13]. In another example, phylogenetic analysis of Aphidiinae parasitoid wasps showed that nearly all species previously categorised as generalists were in fact cryptic, host specialised, species complexes [14]. Even when the occurrence of true generalist species has been demonstrated, a degree of host specialisation may be inevitable. In the generalist oomycete plant pathogen *Albugo candida*, host adapted races suppress plant immunity which facilitates colonisation by non-specialist lineages providing opportunities for gene flow (or genetic introgression) between host races, enabling host range expansion [15]. As such, genuine generalists remain rare, and how such parasites manage to keep up in multilateral coevolutionary arms races remains an evolutionary enigma.

The green peach aphid *Myzus persicae* is an extreme example of a genuine generalist, being able to colonise more than 100 different plant species from 40 plant families [16]. As in many other aphid species, *M. persicae* has a complex life cycle that consists of both sexual and parthenogenetic (clonal) stages. Sexual reproduction occurs in autumn on *Prunus* spp. and produces overwintering eggs from which parthenogenetically reproducing nymphs emerge in the spring [17, 18]. These clonally reproducing individuals soon migrate to an extraordinarily diverse range of secondary host species, including many agriculturally important crop species [19]. In areas where *Prunus* spp. are mostly absent, such as in the United Kingdom, *M. persicae* becomes facultatively asexual, remaining on its secondary hosts all year round [19]. In both cases, clonal populations of *M. persicae* are found on diverse plant species. For example, *M. persicae* clone O populations are found on multiple crop species in the UK and France, including *Brassica* species, potato and tobacco [20, J. C. Simon, pers. Communication].

To investigate the genetic basis of generalism in *M. persicae*, we sequenced the genomes of two *M. persicae* clones, G006 from the USA and O from the UK and the transcriptomes of clone O colonies reared on either *Brassica rapa* or *Nicotiana benthamiana*. These two plant species produce different defence compounds shown to be toxic to insect herbivores [21, 22] presenting distinct challenges to aphid colonisation. Here we provide evidence that the transcriptional adjustments of co-regulated and aphid-expanded multiple member gene families underpin the phenotypic plasticity that enables rapid colonisation of distinct plants by *M. persicae* clone O.

## Results

### M. persicae genome sequencing and annotation

To generate a high quality *M. persicae* genome assembly we sequenced a holocyclic line of the US clone G006 [23] using a combination of Illumina paired end and mate pair libraries (Additional File 1: Table S1). The size of the assembled *M. persicae* genome was 347 Mb including ambiguous bases, representing over 82% of the total genome size as estimated from a kmer analysis of the raw reads (421.6 Mb). The assembly consists of 4,018 scaffolds > 1 kb with an N50 scaffold length of 435 Kb and an average coverage of 51x (Table 1). A total of 18,529 protein-coding genes (30,127 isoforms) were predicted using an annotation workflow incorporating RNA-Seq and protein alignments. We also generated a draft assembly of *M. persicae* clone O, the predominate genotype in the UK [24]. The clone O genome was independently assembled to a size of 355 Mb with 18,433 protein coding genes (30,247 isoforms) annotated, validating the genome size and number genes identified in the G006 assembly (Table 1). Contiguity of the clone O assembly was lower than that of G006, with the assembled genome containing 13,407 scaffolds > 1 Kb and having an N50 scaffold length of 164 Kb. Full details of the assembly, annotation and validation of both genomes are given in Additional File 2.

**Table 1.** Genome assembly and annotation summary.

|  | *M. persicae* |  | *A. pisum* |
| --- | --- | --- | --- |
| **Statistic** | **Clone O** | **Clone G006** | **Release 2.1b** |
| **Genome** |  |  |  |
| # sequences (> = 1 kb) | 13407 | 4018 | 12969 |
| Largest scaffold | 1,018,155 | 2,199,663 | 3,073,041 |
| Total length | 354,698,803 | 347,304,760 | 541,675,471 |
| Total length (> = 1 kb) | 354,698,803 | 347,300,841 | 532,843,107 |
| N50 | 164,460 | 435,781 | 570,863 |
| GC% | 30.19 | 30.03 | 29.69 |
| # Ns | 11,562,637 | 1,836,185 | 36,934,320 |
| Median kmer Coverage | 44X | 51X | NA |
| CEGMA (% complete genes) | 94.35 | 94.35 | 93.15 |
| **Annotation** |  |  |  |
| Gene Count (Coding) | 18,433 | 18,529 | 36,939 |
| Total transcripts | 30,247 | 30,127 | 36,939 |
| Transcripts per gene | 1.64 | 1.63 | 1.00 |
| Transcript mean size cDNA (bp) | 2119.36 | 2163.47 | 1964.11 |

### *Metabolic pathways are similar in* M. persicae *and* A. pisum

A global analysis of the metabolism enzymes of *M. persicae* was generated based on the annotated gene models (Additional File 3) and is available in the ArthropodaCyc metabolic database collection (http://arthropodacyc.cycadsys.org/) (Baa-Puyoulet *et al.*, in press). Metabolic reconstruction in *A. pisum* has highlighted the metabolic complementarity between the aphid and its obligate bacterial symbiont, *Buchnera aphidicola*, with the symbiont generating essential amino acids for the aphid [25]. We compared the amino acid metabolism pathways identified in the two clones of *M. persicae* with those previously identified in *A. pisum*[25, 26]. *A. pisum* and the two *M. persicae gene sets* share 170 enzymes belonging to known amino acid metabolism pathways. *A. pisum* has 22 enzymes that were not found in either of the two *M. persicae* gene sets, and *M. persicae* has 13 enzymes that were not found in *A. pisum*. As previously shown in *A. pisum* the *M. persicae* amino acid metabolism pathways appear complementary with that of *B. aphidicola*. Also, similar to *A. pisum* and *D. noxia* (manual Blast analyses, data not shown), *M. persicae* lacks the tyrosine (Tyr) degradation pathway that is present in all insects included in ArthropodaCyc at the time of writing, indicating that the lack of this pathway may be common feature of aphids. As such, the ability of *M. persicae* to colonise multiple plant species is unlikely to involve specific metabolic pathways that are absent in more specialised aphids.

### Dynamic gene family evolution in aphids

To investigate gene family evolution in aphids and to understand if specific gene repertoires may contribute to *M. persicae* ability to have a broad plant host range, we conducted a comparative analysis of *M. persicae* genes with those of the specialist aphid *A. pisum,* and 19 other arthropod species. Genes were clustered into families based on their protein sequence similarity, inferred from an all-vs-all blastp search, using the Markov Cluster Algorithm (MCL) [27]. Herein, unless otherwise stated, we use the term ‘gene family’ to represent clusters generated by MCL. Phylogenetic relationships amongst the included taxa were inferred using maximum likelihood (ML) with RAxML [28] based on 66 strict, single copy orthologs found in all species (Additional File 4: Figure S1). Relative divergence times were then estimated based on this topology with RelTime [29] (Figure 1). With the exception of the placement of *Pediculus humanus*, all phylogenetic relationships received maximum support and are in agreement with a recently published large-scale phylogenomic study of insects [30]. Annotation of the *M. persicae* genome reveals a gene count approximately half that of the specialist aphid *A. pisum*, and similar to that of other insect species (Figure 1), implying the massive increase in gene content observed in *A. pisum* [26] may not be a general feature of aphid species. Using our comparative dataset, we find that the larger gene count of *A. pisum* compared to *M. persicae* is explained by two features, an increase in lineage specific genes and widespread duplication of genes from conserved families (Figure 1). *A. pisum* has approximately 4 times the number of lineage specific genes than *M. persicae* (8,876 vs. 2,275) and a greater number of genes in families with patchy orthology relationships across insects (5,628 vs. 7,042 respectively). The higher number of broadly conserved genes in *A. pisum* is due to widespread gene duplication rather than differential loss of whole gene families in *M. persicae* with 75% (3,336 / 4,406) of *A. pisum* gene families that have patchy orthology in arthropods also found in *M. persicae*. Furthermore, the mean size of these families has increased by 82% in *A. pisum* (3.55 vs. 1.95, Mann–Whitney *U* p < 0.00005). This is underlined by the pattern across all genes, with *A. pisum* having a significantly higher proportion of multi-copy genes than *M. persicae* (23,577 / 36,193 in *A. pisum* vs. 9,331 / 18,529 in *M. persicae*, Chi square test: χ^2^=1220.61, d.f.=1, p = 2.02 ×10^−267^).

**Figure 1:**
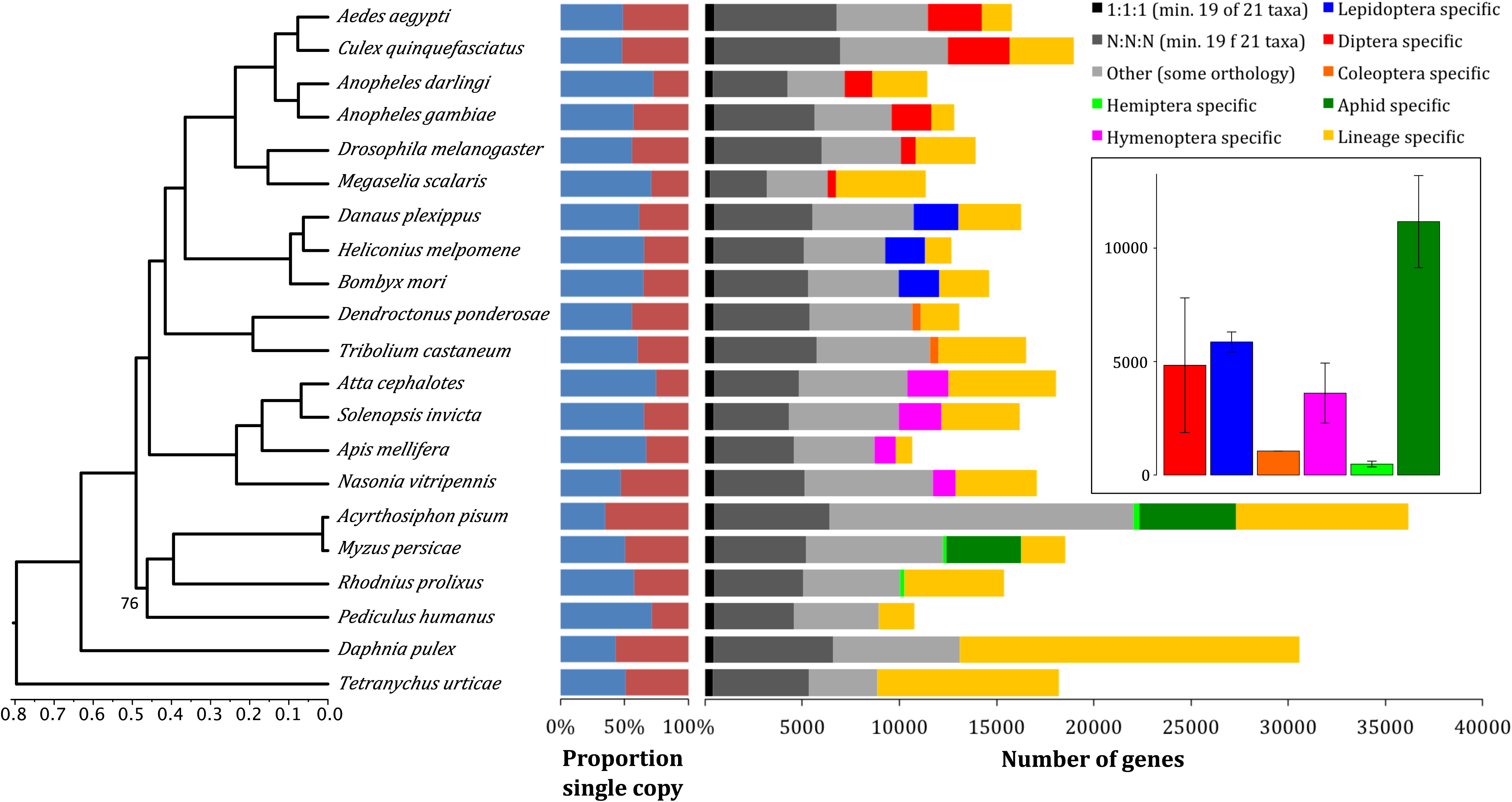
High rate of lineage specific gene accumulation in aphids relative to all other insect orders. Figures show arthropod phylogenetic relationships, per genome proportions of single copy (blue) and duplicated (red) genes, and orthology relationships amongst arthropod genes based on gene family clustering with MCL [27]. Phylogenetic relationships amongst arthropod species included for gene family clustering 27were estimated using RAxML [28] based on a protein alignment of 66 single copy orthologs found in all taxa. This topology and protein alignment was then used to infer relative divergence times with RelTime [29] under an LG substitution model.27 Inset shows relative rate of lineage specific gene accumulation for all included insect orders and comparison with aphids. Error bars show standard deviation of species within a given grouping. Relative rates of lineage specific gene accumulation were calculated for each species by dividing the number of group specific genes (either order specific or aphid specific) by the crown plus stem age for the given group (in relative divergence time).

In addition to the differences observed between the two aphid species, there also appears to have been considerable change in gene content during aphid evolution relative to other insect orders. After accounting for evolutionary divergence, the rate of accumulation of aphid-specific genes is higher than the accumulation of lineage-specific content in any other insect order (Figure 1). GO term enrichment analysis of these genes shows they are enriched for biological processes including detection and response to chemical stimuli, metabolic regulation and regulation of transcription, processes likely important in aphid evolution and diversification (Additional File 5: Figure S2 and Additional File 6: Table S2).

Modelling of gene gain and loss in widespread gene families across the arthropod phylogeny also highlights the dynamic pattern of gene family evolution in aphids (Additional File 7: Figure S3). After correcting for evolutionary distance between species, *A. pisum* has the highest rate of gene family expansion of any arthropod species (Additional File 7: Figure S3). *M. persicae* has also undergone a relatively high number of gene family expansions over a short period of time compared to other arthropod species, but has significantly fewer expanded gene families than *A. pisum* (114 / 4983 vs. 538 / 4983; Chi square test: χ^2^= 295.03, d.f.=1, p= 3.984 ×10^−66^), and overall it has undergone a net decrease in gene family size. As such, gene gain in *M. persicae* appears to be restricted to a smaller subset of gene families than in *A. pisum*. This was also confirmed using a more inclusive set of gene families (6,148 families found in both aphids as well as at least one other species) with a binomial test to identify significant expansion (173 / 6148 vs. 391 / 6148; Chi square test: χ^2^= 88.31, d.f.=1, p= 5.59 ×10^−21^). Interestingly, 85 % of gene family expansions in *M. persicae* were shared with *A. pisum*. This suggests that a subset of *M. persicae* gene families may have been selected to retain high ancestral copy number, or have experienced parallel, lineage-specific, duplication, against a background of reduced expansion genome wide. Full details of all expanded families are given in (Additional File 8: Table S3).

### Genome streamlining in a generalist aphid

Differences in overall gene count and patterns of gene family evolution between *M. persicae* and *A. pisum* may be the result of a shift in gene duplication rate, altered selective regimes acting on duplicate retention (i.e. genome streamlining), or a combination of the two. To test this we conducted a synonymous (*d*_*S*_) and non-synonymous (*d*_*N*_) substitution rate analysis and found evidence of increased genome streamlining in the generalist aphid *M. persicae* (Figure 2A and Additional File 9: Figure S4). The age distribution of paralogs in *M. persicae* and *A. pisum* shows that gene duplicates have accumulated steadily in both species with a continuing high rate of duplication (Figure 2A). However, we observe marked differences in the retention rates of ancestrally duplicated genes between the two species. Using average *d_S_* between *M. persicae* and *A. pisum* 1:1 orthologs (*d_S_*=0.26) as a cut off to identify ancestral (pre-speciation) duplicates, we find a significantly greater loss rate in *M. persicae* than *A. pisum*. In *A. pisum*, we found 382 genes that duplicated before speciation, and of those, *M. persicae* has lost one or both paralogs in 224 families, (59% loss). We detected 285 families that duplicated before speciation in *M. persicae*, and of those, 69 families lost one or both paralogs in *A. pisum* (24% loss) (Chi-square test: χ^2^=78.55, d.f.=1, p=7.82 ×10^−19^). Consistent with genome streamlining, we also observe stronger purifying selection in ancestral duplicates retained in *M. persicae* than in *A. pisum* (Figure 2B and Additional File 9: Figure S4).

**Figure 2:**
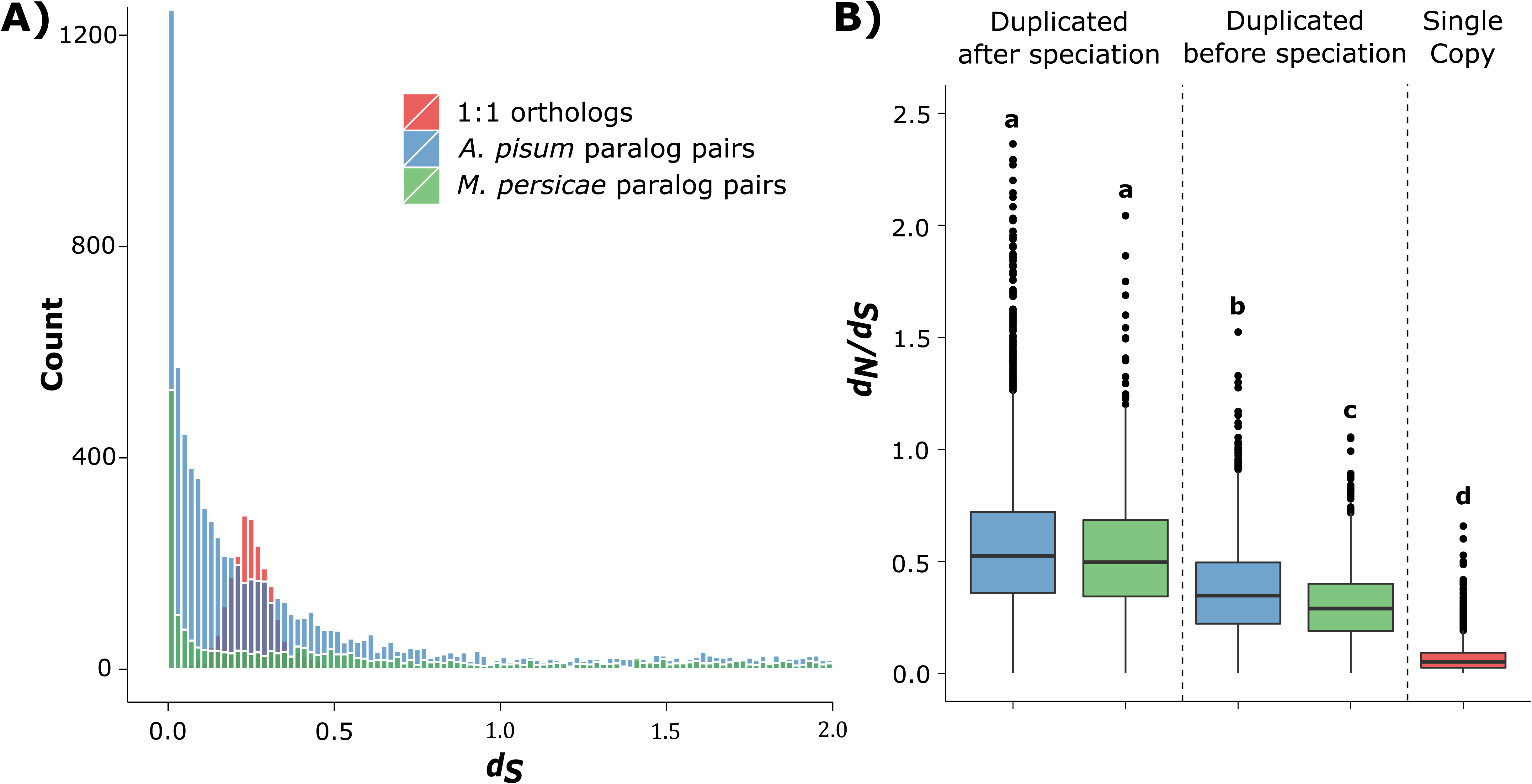
*M. persicae* experienced greater gene loss rates (**A**) and stronger purifying selection in retained ancestral duplicates (**B**) than *A. pisum*. **A)** Age distribution of duplicated genes in *M. persicae* and *A. pisum*. The number of synonymous substitutions per synonymous site (*d_S_*) was calculated between paralog pairs for *M. persicae* (green) and *A. pisum* (blue) using the YN00 [87] model in PAML [78]. For each duplicated gene only the most recent paralog was compared. Pairwise *d_S_* was also calculated for 1:1 orthologs between *M. persicae* and *A. pisum* (red), the peak in which corresponds to the time of speciation between the two aphid species. After filtering, 1,955 *M. persicae* paralog pairs, 7,253 *A. pisum* paralog pairs and 2,123 1:1 orthologs were included for comparison. Mean *d_S_* of 1:1 orthologs between *A. pisum* and *M. persicae* was 0.26. **B)** Box plots showing median *d_N_*/*d_S_* for *A. pisum* and *M. persicae* paralog pairs that duplicated before and after speciation of the two aphid species and for 1:1 orthologs between the two species. Older duplicate genes have lower *d*_*N*_/*d_S_* than recently duplicated genes (since speciation) indicating stronger purifying selection in ancestral versus recent duplicates. Additionally, older duplicate genes in *M. persicae* have significantly lower *d_N_*/*d_S_* than in *A. pisum* (Mann-Whitney U = 1816258, *M. persicae*: 1,348 paralog pairs, *A. pisum*: 3,286 paralog pairs, p = < 0.00001) indicating stronger genome streamlining in *M. persicae* than in *A. pisum*.. Box plots are shaded by species as in **A**.

### A phylome resource for aphids

A phylome resource (the complete collection of gene trees) for *M. persicae* and 259 all taxa included in the comparative analysis was also generated, and is available for download or to browse at PhylomeDB [31]. Gene trees were scanned to infer duplications and speciation events and to derive orthology and paralogy relationships among homologous genes [32]. Duplication events were assigned to phylogenetic levels based on a phylostratigraphic approach [33] and duplication densities calculated on the branches of the species tree leading to *M. persicae*. In agreement with the comparative analysis above, a high rate of duplication was observed on the branch leading to *M. persicae* and *A. pisum* and relatively low rate of duplication observed in *M. persicae* (for full methods and results see Additional File 10).

### *Host transition in* M. persicae *involves transcriptional plasticity of aphid specific and aphid expanded genes that constitute gene clusters in the aphid genome.*

In order to examine how genetically (near) identical *M. persicae* clones are able to colonie divergent host species, clone O colonies were started from single females and reared on *Brassica rapa* (Chinese cabbage, Brassicaceae), and subsequently transferred to *Nicotiana benthamiana* (Solanaceae). The two clonally reproducing populations were reared in parallel on these plants for one year and their transcriptomes sequenced. Comparison of these transcriptomes identified 171 differentially expressed (DE) genes putatively involved in host adjustment (DEseq, > 1.5 fold change, 10% false discovery rate (FDR); Figure 3; Additional File 11: Table S4).

**Figure 3:**
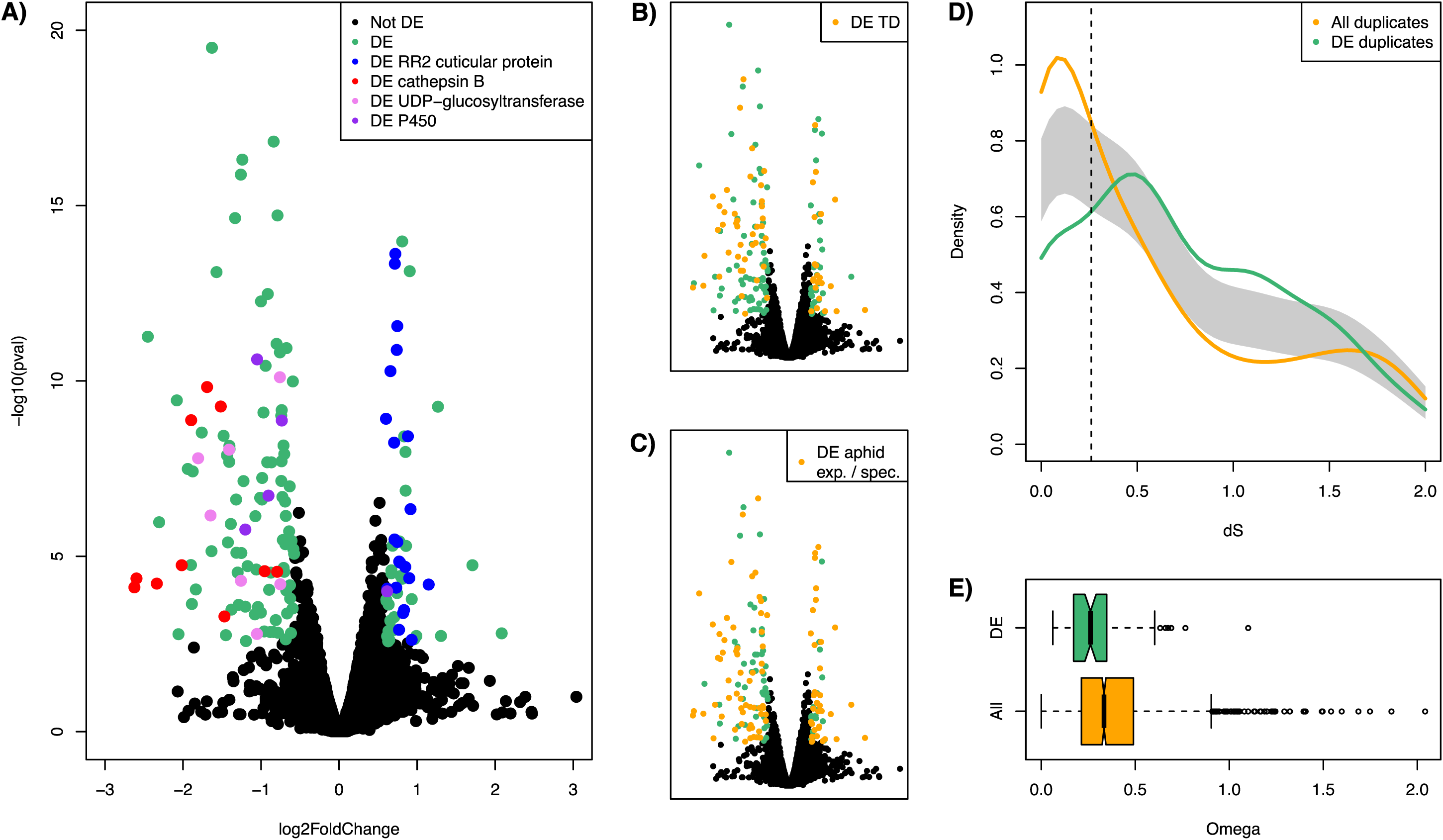
The set of differentially expressed genes of *M. persicae* clone O reared on *B. rapa* and *N. benthamiana* is enriched for (**A**) genes belonging to gene families with known functions, (**B**) tandemly duplicated genes in the *M. persicae* genome, (**C**) genes belonging to gene families expanded in aphids or unique to aphids, (**D**) duplicated genes before *M. persicae* and *A. pisum* diverged and (**E**) genes with stronger purifying selection than the genome wide average. **A - C)** Volcano plots of differentially expressed genes of *M. persicae* reared on *B. rapa* and *N. benthamiana*. Negative log_2_ fold changes indicate up-regulation on *B. rapa* and positive values indicate up-regulation on *N. benthamiana* **A)** Differentially expressed genes from four gene families that have the highest number of differentially expressed genes are highlighted. These are: RR2 cuticular proteins (n=22), Cathepsin B (n=10), UDP-glucosyltransferase (n=8) and Cytochrome P450 (n=5). **B)** The set of differentially expressed genes is enriched for tandemly duplicated genes. **C)** The set of differentially expressed genes is enriched for genes from families that are either significantly expanded in aphids compared to other arthropods (binomial test, main text), or are unique to aphids. **D)** Time since most recent duplication (measured as *d_S_*) for all paralogs in the *M. persicae* genome compared to those differentially expressed upon host transfer. Duplicated genes implicated in host adjustment (at least one of the pair differentially expressed) have a significantly different distribution to the genome wide average (p < 0.05, permutation test of equality) and are enriched for genes that duplicated before *M. persicae* and *A. pisum* diverged. **E)** *d_N_*/*d_S_* distribution for duplicated genes differentially expressed upon host transfer vs. the genome wide average. Duplicated genes involved in host adjustment are under significantly stronger purifying selection than the genome wide average (median *d_N_*/*d_S_* = 0.2618 vs. 0.3338, Mann–Whitney *U*= 105,470, p = 1.47 × 10^−4^, two-tailed).

The set of differentially expressed genes was significantly enriched for genes from multigene families compared to the genome as a whole (126 / 171 DE vs. 9,331 / 18,529 genome wide (GW), Chi square test: χ^2^= 36,88, d.f.=1, p = 6.92 × 10^−10^; Figure 3A, C). Furthermore, many of the differentially expressed genes are from aphid expanded or aphid specific gene families (105 / 171 DE vs. 3,585 / 18,529 GW, Chi square test: χ^2^= 195.62, d.f.=1, p = 1.89 × 10^−44^; Figure 3C), highlighting the important role of aphid genomic novelty in *M. persicae* colonisation of diverse plant species. In most cases, gene families were uni-directionally regulated with 64 families up-regulated on *B. rapa* and 36 families up-regulated on *N. benthamiana* (Additional File 11: Table S4). Genes from only 6 families were bi-directionally regulated on the plant hosts. Of these, multiple genes of the UDP-glycosyltransferases, maltase-like, P450 monooxygenases and facilitated trehalose transporter Tret1-like were up-regulated on *B. rapa* and single genes in each of these families on *N. benthamiana* (Additional File 11: Table S4).

The cathepsin B and Rebers and Riddiford subgroup 2 (RR-2) cuticular protein [34] families, which have the highest number genes differentially expressed upon host transfer (Figure 3A), typify the way *M. persicae* gene families respond to host transfer. Members of these families are uni-directionally regulated, with Cathepsin B genes up-regulated in aphids reared on *B. rapa* and RR-2 cuticular proteins up-regulated in aphids reared on *N. benthamiana.* Further annotation of the cathepsin B and RR-2 cuticular protein genes (Additional File 3) and phylogenetic analyses of these genes with other hemipteran species reveals that differentially expressed genes from these families cluster together in aphid expanded, and in the case of cathepsin B, *M. persicae* expanded, clades (Figure 4A and Additional File 12: Figure S5A) for full methods and results see Additional File 10]. We also found that cathepsin B and RR-2 cuticular proteins regulated in response to host change are clustered together in the *M. persicae* genome with differentially expressed members forming tandem arrays within scaffolds (Figure 4B and Additional File 12: Figure S5B). Differentially expressed UDP-glycosyltransferase, P450 monooxygenases and lipase-like are also arranged as tandem repeats (Additional Files 13 – 15: Figures S6 – S8), and more generally, tandemly duplicated genes were overrepresented among the differentially expressed genes (65 / 171 DE vs. 1111 / 18529 GW, Chi-square test, χ^2^= 314.66, d.f.= 1, p = 2.10 × 10^−70^; Figure 3B) highlighting the tendency of genes regulated in response to host change to be clustered in the *M. persicae* genome.

**Figure 4:**
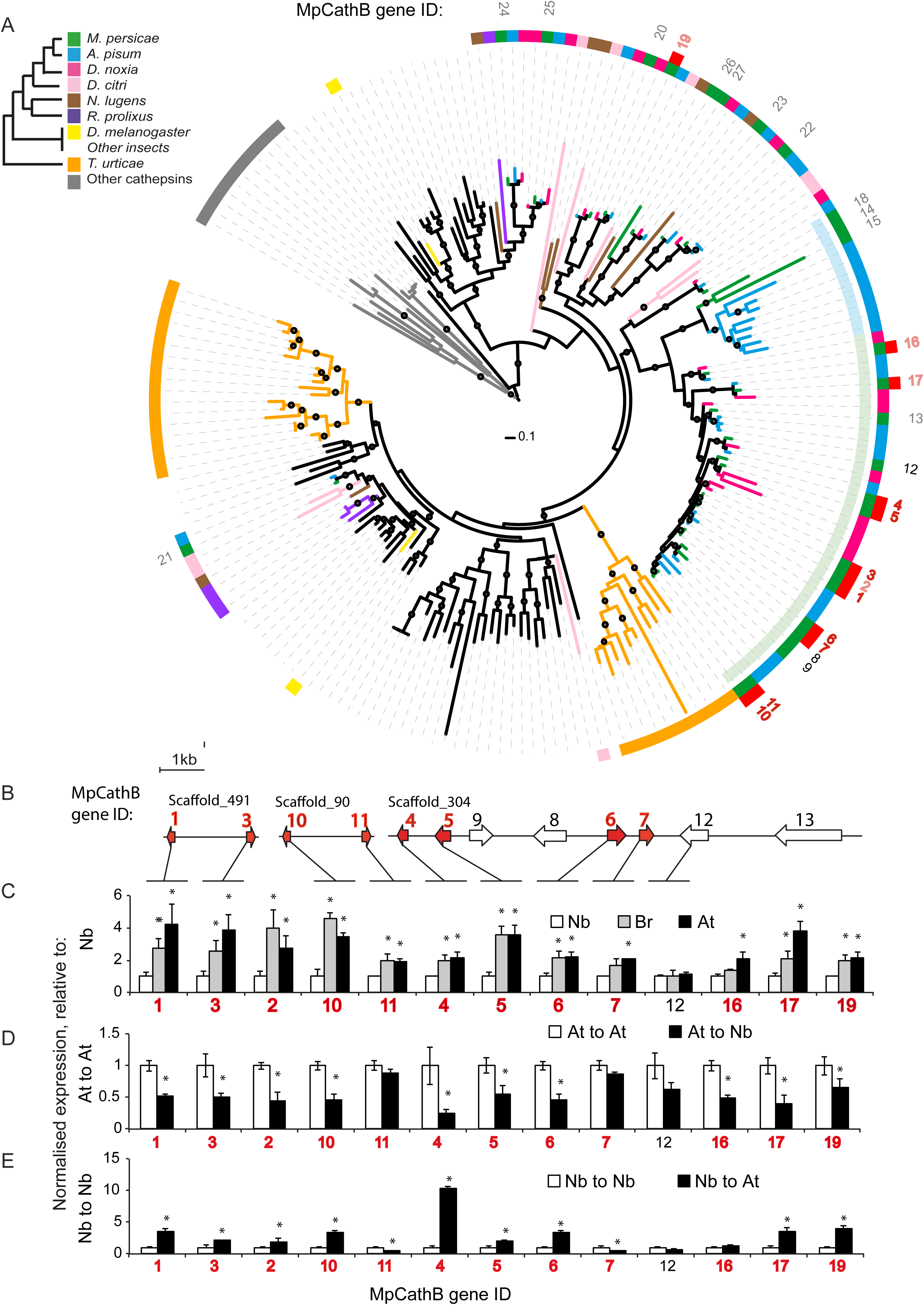
Cathepsin B genes that are differentially expressed upon *M. persicae* host change belong predominantly to a single aphid-expanded clade and form gene clusters in the *M. persicae* genome. **A)** Maximum likelihood phylogenic tree of arthropod cathepsin B protein sequences. The sequences were aligned with Muscle [72] and the phylogeny estimated using FastTree [88] (JTT + CAT rate variation). Circles on branches indicate SH-like local support values >80%, scale bar below indicates 0.1 substitutions per site. Rings from outside to inside: ring 1, *M. persicae* cathepsin B (MpCathB) gene identities (IDs) with numbers in red indicating up-regulation of these genes in *M. persicae* reared for one year on *B. rapa* relative to those reared for one year on *N. benthamiana*, and bold font indicating location on the cathepsin B multigene clusters shown in **B**; ring 2, red squares indicating MpCathB genes that are differentially expressed upon *M. persicae* host change; ring 3, cathB genes from different arthropods following the colour scheme of the legend in the upper left corner and matching the colours of the branches of the phylogenetic tree; ring 4, aphid expanded (AE) clades with AE_Clade I labelled light green and AE_Clade II light blue. **B)** MpCathB multigene clusters of the *M. persicae* genome. Lines indicate the genomic scaffolds on which the MpCathB genes are indicated with block arrows. Gene IDs above the genes match those of the phylogenetic tree in A, with block arrows and fonts highlighted in red being differentially expressed upon host change. Scale bar on right shows 1 kb.**C)** Relative expression levels of MpCathB genes of *M. persicae* at 7 weeks being reared on *N. benthamiana* (Nb), *B. rapa* (Br) and *A. thaliana* (At). Numbers under the graphs indicate MpCathB gene IDs with those in red font DE as in A. Batches of five adult females were harvested for RNA extraction and qRT-PCR assays. Bars represent expression values (mean ± standard deviation (SD)) of three independent biological replicates. \**p*<0.05 (Anova with Fishers LSD to control for multiple tests). **D)** As in C, except that individual aphids reared on At were transferred to At (At to At) or Nb (At to Nb) and harvested at 2 days upon transfer. **E**) As in D, except that individual aphids reared on Nb were transferred to Nb (Nb to Nb) or At (Nb to At) and harvested at 2 days upon transfer.

In many parasites recent, lineage-specific, gene family expansions have been implicated in host range expansion and transitions to generalism, for example in the nematode genus *Strongyloides*[35] and the ascomycete genus *Metarhizium*[36]. We therefore tested for the presence of recently duplicated genes involved in *M. persicae* host colonisation (differentially expressed on host transfer) by estimating the coalescence times of these genes and comparing them to the aphid phylogeny. Contrary to our expectations, the analysis of pairwise substitution patterns between duplicated differentially expressed genes and their closest paralog show that these genes are older than the genome wide average, with the differentially expressed gene set enriched for gene duplicates that arose before the divergence of *M. persicae* and *A. pisum* (paralog pairs *d_S_* 0.26 − 2.00: DE duplicated = 75 / 97, whole genome = 1,348 / 2,414, Chi square test: χ^2^= 15.87, d.f.=1, p = 6.79 × 10^−5^) (Figure 3D). In addition, we found that host regulated genes appear to be under stronger purifying selection than the genome wide average with paralog pairs containing at least 1 differentially expressed gene having median *d_N_*/*d_S_* significantly lower than for all paralog pairs in the genome (median *d_N_*/*d_S_* = 0.2618 vs. 0.3338, Mann–Whitney *U* = 105,470, p = 1.47 × 10^−4^) (Figure 3E, Additional File 16: Table S5). This suggests that most of the genetic variation utilised during host colonisation was present in the common ancestor of the two aphid species, and hence *Myzus* specific gene duplication alone does not represent the evolutionary innovation that enables a generalist lifestyle. Rather, generalism could be facilitated by the plastic expression of predominantly pre-existing genetic variation - in this instance, aphid specific gene duplicates.

### Gene expression changes upon host transfer occur rapidly

To further investigate gene expression plasticity in *M. persicae* upon transfer to diverged hosts, we investigated differential gene expression of aphids transferred from *B. rapa* to *N. benthamiana* and allowed adjustment on their new hosts for 7 weeks, this time also including a transfer from *B. rapa* to *Arabidopsis thaliana*. *M. persicae* clone O successfully colonised all three host species with no significant differences observed between development time, reproduction rate, longevity or weight (Additional File 17: Figure S8.5). We used cathepsin B and RR-2 cuticular proteins identified as differentially expressed by RNA-Seq as marker genes and measured their expression by qRT-PCR. All differentially expressed cathepsin B and RR-2 cuticular protein genes for which specific primers could be designed (the majority) found to be differentially expressed in the RNA-Seq experiments were also differentially expressed in the qRT-PCR experiments. Furthermore, we find similar expression patterns for aphids reared on Brassicaceae species with cathepsin B copies up-regulated on *B. rapa* and *A. thaliana* relative to *N. benthamiana* (Figure 4C) and RR-2 cuticular proteins down-regulated (Additional File 12: Figure S5C).

To investigate how fast gene expression changes upon host transfer, individual aphids (3-day old nymphs) were transferred from *A. thaliana* to *N. benthamiana* and vice versa, or to the same host, and expression of cathepsin B and RR-2 cuticular protein genes measured after two days by qRT-PCR. Cathepsin B gene expression went up in aphids transferred from *N. benthamiana* to *A. thaliana* and down in aphids transferred from *A. thaliana* to *N. benthamiana* (Figure 4D,E). Conversely, expression of RR-2 cuticular protein genes went down in aphids transferred from *N. benthamiana* to *A. thaliana* and up in aphids transferred from *A. thaliana* to *N. benthamiana* (Additional File 12: Figure S5D,E). No significant change was observed when aphids were transferred to the same plant species (from *A. thaliana* to *A. thaliana*, or *N. benthamiana* to *N. benthamiana*). Hence, expression levels of cathepsin B and RR-2 cuticular protein genes adjust quickly upon host change (within 2 days) and regulated in a coherent, host dependent, fashion.

### *Cathepsin B contribute to* M. persicae *fitness in a plant host dependent manner*

To test whether targets of transcriptional plasticity in *M. persicae* have direct fitness affects we conducted plant-mediated RNAi knockdown [37, 38] of cathepsin B genes identified as differentially expressed upon host transfer. We focused on cathepsin B as the majority (11 out of 12) of gene copies differentially expressed upon host transfer are located in a single, *M. persicae* expanded clade (Cath_Clade I) of the cathepsin B phylogeny (Figure 4A) and have 69-99% nucleotide sequence identities to one-another (Additional File 18). As such, a single dsRNA construct can be used to knock down multiple cathepsin B genes. In contrast, the clade containing the majority of differentially regulated RR-2 cuticular protein genes is larger and more diverse (Additional File 12: Figure S5), presenting a challenge for using the RNAi-mediated approach to examine how these genes act together to enable *M. persicae* colonisation. Three independent stable transgenic *A. thaliana* lines producing dsRNAs targeting multiple cathepsin B genes (At_dsCathB 5-1, 17-5 and 18-2; Additional File 18) were generated. The expression levels of all Cath_Clade I genes except MpCath12 were down-regulated in *M. persicae* reared on these lines (Figure 5A) in agreement with MpCath12 having the lowest identity to the dsRNA sequence (73% vs > 77% for other copies) (Additional File 18). Aphids on the three At_dsCathB lines produced about 25% fewer progeny (p<0.05) compared to those reared on the At_dsGFP control plants (Figure 5B) indicating that the cathepsin B genes contribute to *M. persicae* ability to colonise *A. thaliana*.

**Figure 5:**
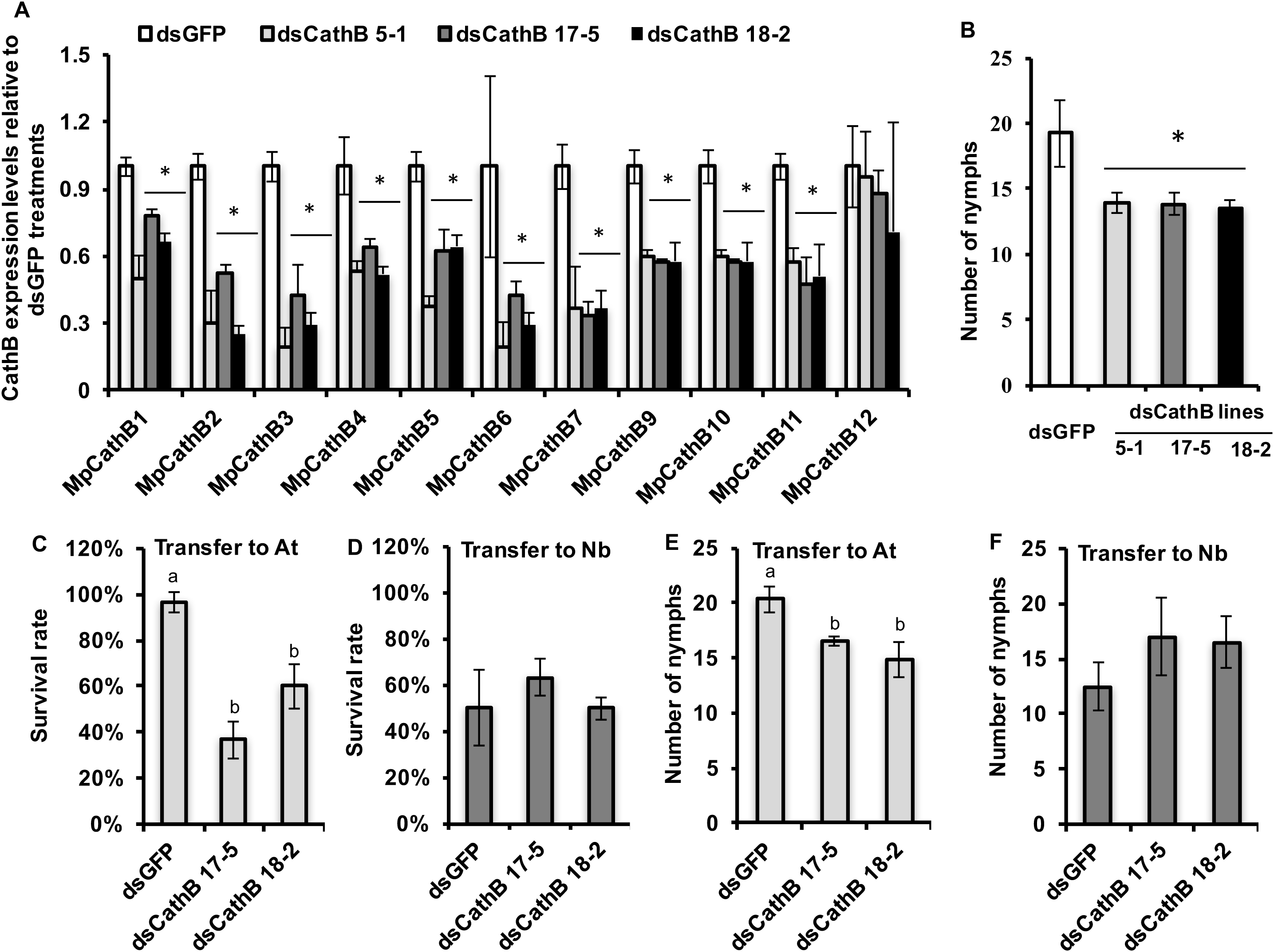
RNA interference (RNAi)-mediated knock-down of the expression of multiple cathepsin B genes reduces *M. persicae* survival and fecundity on *A. thaliana*. **A)** Relative cathepsin B (CathB) expression levels (compared to aphids on *dsGFP* (control) plants) of *M. persicae* on three independent transgenic lines (line 5-1, line 17-5, and line 18-2) producing double-stranded (ds) RNA corresponding to multiple *M. persicae* cathepsin B genes (dsCathB) (Figure 3A, Additional File 19: Figure S9). Aphids were reared on the transgenic lines for four generations. Batches of five adult females were harvested for RNA extraction and qRT-PCR assays. Bars represent expression values (mean ± standard deviation (SD)) of three independent biological replicates. **B)** CathB-RNAi *M. persicae* produces less progeny compared to control (dsGFP-treated) aphids on *A. thaliana*. Five nymphs were transferred to single plants and produced nymphs on approximately day 5. Nymph counts were conducted on days 7, 9, and 11 and removed. Columns show the mean ± SD of the total nymph counts for these three days of three biological replicates, with each replicate consisting nymphs produced by 15 aphids at 5 aphids per plant (n = 3 plants)**. C and D)** Survival rates of CathB-RNAi and control (dsGFP-exposed) *M persicae* on non-transgenic *A. thaliana* (At) and *N. benthamiana*^(Nb) plants. Ten 3^rd instar nymphs on *dsCathB* and *dsGFP* transgenic plants were transferred to non-transgenic plants, survival rate were recorded two days later. Bars represent mean ± SD of three biological replicates, with each replicate consisting of the survival rates of 30 aphids at 10 aphids per plants (n = 3 plants). **E and F)** Fecundity rates of CathB-RNAi and control (dsGFP-exposed) *M persicae* on non-transgenic *A. thaliana* (At) and *N. benthamiana* (Nb) plants. Nymph counts were conducted as in **B**. Asterisks (*) and different letters (a, b) above the bars indicate significant difference at *p*<0.05 (Anova with Fishers LSD to control for multiple tests).

To examine the impact of cathepsin B on the ability of *M. persicae* to adjust to host change, the cathB-RNAi aphids were transferred from At_dsCathB lines to non-transgenic *A. thaliana* and *N. benthamiana* plants and examined for survival and fecundity. In agreement with previous data [38], we found that the genes targeted by RNAi remain down regulated at 2 days upon transfer from At_dsCathB lines to non-transgenic plants (Additional File 19: Figure S9). Upon transfer to *A. thaliana*, the cathB-RNAi aphids had lower survival and reproduction rates than the dsGFP-exposed (control) aphids (Figure 5C,E). In contrast, no decline in survival and reproduction was seen of the cathB-RNAi aphids compared to the dsGFP-exposed aphids upon transfer to *N. benthamiana* (Figure 5D,F). Thus, cathB knock down impacts *M. persicae* fitness differentially depending on the host plant species. Together these data provide evidence that adjustment of the cathepsin B gene expression levels between *A. thaliana* and *N. benthamiana* contributes to the ability of *M. persicae* to colonise both plant species.

## Discussion

So far, genomic studies of polyphagy and generalism have primarily focused on genetic adaptation and have led to the identification of specific genetic elements that are present in the genomes of one race (or biotype) versus another and that enable these races to be host-specific [13, 15, 26]. In such cases, whilst the species as a whole may be considered polyphagous, individuals are not. Here, we have investigated the genome and transcriptome of the genuine generalist *Myzus persicae*. We demonstrate the striking ability of *M. persicae* to colonise divergent host plant species by conducting host transfer experiments using individuals from a single, clonally reproducing line (Clone O), and allowing them to adjust to three distinct host plant species from two plant families. We show that generalism in *M. persicae* is associated with rapid transcriptional plasticity of often aphid-specific gene copies from multi-gene families that are uni-directionally regulated. Furthermore, we show that disrupting the transcriptional adjustment of a gene family with high levels of differentially expressed upon host transfer (cathepsin B), using plant mediated RNAi, has host dependent fitness costs for *M. persicae*, suggesting that host associated transcriptional plasticity is adaptive in *M. persicae*.

Contrary to expectations, the majority of genes differentially regulated upon host transfer originate from ancestral aphid duplication events rather than more recent lineage-specific duplications. Additionally, comparative analysis of all *M. persicae* gene families with other arthropods showed that, whilst gene family evolution appears to have been highly dynamic during aphid diversification, *M. persicae* does not exhibit widespread gene duplication on the scale of the legume specialist *A. pisum*. This is surprising given that other studies have shown a key role for lineage-specific gene duplication in parasite host range expansions [35, 36]. Although not extensive, recent gene duplication may still play a role in *M. persicae* host adaptation given that some gene families have undergone *M. persicae* specific gene duplication against a background of reduced gene family expansion genome wide. For example, the cathepsin B and UGT gene families have undergone *M. persicae* specific gene duplication and are implicated in host adjustment. These observations are consistent with genome streamlining in *M. persicae,* with functionally important gene duplicates preferentially retained. It therefore seems likely that functionally important lineage-specific gene duplication combined with rapid transcriptional plasticity of a broader, aphid-specific gene repertoire, consisting of selectively retained gene duplicates, underpins the generalist feeding habit in *M. persicae*.

Transcriptional plasticity has also been implicated in host adjustment in generalist spider mite and butterfly species [39, 40]. This suggests a key role for transcriptional plasticity in plant feeding arthropods that have evolved genuine generalism as opposed to cryptic sub-structuring of genetic variation by host species. The mechanisms by which this transcriptional plasticity is achieved are, as yet, unknown. However, given that in *M. persicae* differences in gene expression occur rapidly upon host transfer, and in the absence of genetic variation between host-adjusted lineages (experiments were performed with single aphids in the 2-day transfer experiments and with clonally reproducing individuals derived from a single parthenogenetic female in the 7-week and one-year aphid colonies), epigenetic mechanisms of gene expression regulation are likely responsible. Full length copies of the DNA methyltransferase (DNMT) genes DNMT1a, DNMT1b, DNTM2, DNMT3a and DNMT3b and all components of the histone modification system are present in *M. persicae*, as is the case for other aphid species [41, 42, 43], and epigenetic mechanisms have been shown to regulate plastic traits such as hymenopteran caste-specific behaviour [44].

Genes belonging to aphid-expanded clades of the cathepsin B and RR-2 cuticular protein gene families contribute the largest percentages of differentially regulated genes upon host transfer and are therefore likely to play a key role in the ability of *M. persicae* to colonise members of Brassicaceae and Solanaceae. Cathepsin B proteins may serve digestive functions [45, 46], but are also known virulence factors, as they play major roles in invasion and intracellular survival of a number of pathogenic parasites [47, 48, 49]. For example, RNAi-mediated knock down of *Trypanosoma brucei* cathepsin B leads to clearance of parasites from the bloodstream and prevents lethal infection in mice [50]. In the social aphid *Tuberaphis styraci*, cathepsin B has been detected as a major component of the venom produced by soldier aphids which is expelled through the stylets and injected into potential predators [51]. In *M. persicae*, three of the differentially expressed cathepsin B genes encode proteins with signal peptides, are expressed in the *M. persicae* salivary gland [23] and peptides corresponding to cathepsin B are found in proteome analyses of *M. persicae* saliva [52], suggesting they come into direct contact with plant components during feeding. Interestingly, cathepsin B genes involved in host adjustment have functionally diverged in *M. persicae* relative to other aphid species. Most of the differentially expressed cathepsin B genes belong to Cath_Clade_I, which has expanded in *M. persicae* relative to *A. pisum* and *D. noxia* (Figure 4A). Functional analysis of genes in this clade shows that most *M. persicae* copies possess a complete cysteine peptidase domain consisting of a propeptide domain and both cysteine and histidine active sites. In contrast, most *A. pisum* and *D. noxia* copies have an incomplete cysteine peptidase domain (Additional File 20: Figure S10). This is in agreement with previous observations that cathepsin B genes are under selection in aphids [53]. Our finding that cathepsin B genes are differentially regulated in response to *M. persicae* host transfer and that knock down of functionally diverged differentially expressed cathepsin B copies directly impacts *M. persicae* fitness in a host dependent manner highlights the key role of this gene family in aphid evolution.

Cuticular proteins bind chitin via extended version of the RR-1 and RR-2 486 consensus sequences and provide the cuticle with structural support, mechanical protection and mobility [54]. Cuticular protein genes have different expression profiles depending on the insect body part, mechanical property needs, developmental stage, temperature and seasonal photoperiodism [55, 56, 57, 58]. RR-1 proteins are associated mostly with soft and flexible cuticle and RR-2 proteins in hard and rigid cuticles [59, 60]. Interestingly, members of the differentially regulated RR-2 cuticular proteins of *M. persicae* on different plant hosts have identical sequences as those shown to be associated with the acrostyle at the tip (last few microns) of the maxillary stylets of the *M. persicae* mouthparts where the food canal and salivary canals are fused [61]. The acrostyle is in the part of the stylet that performs intracellular punctures during probing and phloem feeding [62] and has a high concentration of cuticular proteins. It also interacts with virus particles that are transmitted by *M. persicae* [61]. Moreover, it is in direct contact with (effector) proteins of the aphid saliva and the plant cell contents, including the phloem sap [62]. Therefore, it is possible that the differential regulation of RR-2 cuticular protein genes enables *M. persicae* to adjust to the different physical and chemical attributes of cell walls, their contents, and defence responses of the diverged plant species.

## Conclusions

We found that *M. persicae* adjustment to diverged plant species involves the 506 unidirectional co-regulation of multigene families that lie within distinct multi-gene clusters in the aphid genome. Differential expression occurs rapidly, within 2 days, indicating strict regulatory control of these gene clusters. Up-regulation of these genes enables *M. persicae* survival and fecundity on the new host. Taken together, this study of the genome sequence of *M. persicae*, comparative genome analyses and experimental study of host change have elucidated specific genes that are involved in the ability of *M. persicae* to colonise members of the Brassicaceae and has provided evidence that the rapid transcriptional plasticity of *M. persicae* plays a role in this aphids ability to adjust to diverged plant species.

## Materials and Methods

### *Preparation of* M. persicae *clones G006 and O for genome sequencing*

Clone G006 was collected from pepper in Geneva, NY, USA in 2003 [23]. Since time of collection, G006 has been maintained on *Brassica oleracea* var. Wisconsin golden acre seedlings in a growth chamber under long day conditions of 16h light: 8 hours of darkness at 20 °C constant temperature in the laboratory of Alexandra Wilson, University of Miami. Clone O is found on multiple crop and weed species in the UK and France, including *Brassica* species, potato and *Nicotiana* species [20, Simon, pers. Communication] and is being reared on Chinese cabbage ( *Brassica rapa;* Brassicaceae), *Arabidopsis thaliana* (Brassicaceae) and *Nicotiana benthamiana* (Solanaceae) in our laboratory. A colony of *M. persicae* clone O starting from a single female was established on *B. rapa* in a growth chamber (14h light, 10h dark at constant 20 °C, 75% humidity) in 2010.

### Genome sequencing

A single paired-end library and two mate-pair libraries were constructed for the G006 clone with insert sizes of approximately 200 (S6), 2000 (S8 MPB) and 5000 (S7 MPA) bp and sequenced with 100bp paired-end run metrics using a version 3 Illumina Hi-Seq paired-end flow cell to give ∼95 Gb of sequencing reads. Illumina library construction and sequencing for clone G006 was performed at the University of Miami’s Center for Genome Sequencing Core at the Hussman Institute for Human Genomics.

For the Clone O genome, three libraries were constructed, two paired-end libraries with an average fragment size of 380 (LIB1672) and 180 (LIB1673) bp and for scaffolding a mate-pair library with an average 8000bp insert size (LIB1472). Libraries were prepared at the Earlham Institute (Norwich, UK) using the Illumina TruSeq DNA Sample Preparation Kit. The resulting DNA libraries were sequenced with 100bp paired-end run metrics on a single lane of an Illumina HiSeq2000 Sequencing System according to manufacturer’s instructions.

### Transcriptome sequencing

Total RNA extracted from *Myzus* G006, tissues include whole female insects (WI), bacteriocytes (dissected from 300 adults) and guts (dissected from 300 adults). All RNA was treated with DNaseI before sending for sequencing at the University of Miami’s Center for Genome Sequencing Core at the Hussman Institute for Human Genomics. Each sample was prepared for mRNA sequencing using an Epicenter PolyA ScriptSeqV2 kit and for small RNA sequencing using an Illumina TruSeq smRNA kit. All sequencing was performed as 2×100 reads on a HiSeq 2000. Each sample was prepared for both mRNA and small RNA sequencing.

To identify genes involved in *M. persicae* host adjustment we sequenced the transcriptomes of clone O colonies reared on *Brassica rapa* and *Nicotiana benthamiana*. Colonies were established from a single asexual female and reared under long-day conditions (14h L 10h D) and constant 20° C and allowed to adapt for one year. Adult asexual females (1-week old) were then harvested in pools of approximately 50 individuals. Three independent pools were harvested from each plant species and RNA extracted using Tri-reagent (Sigma) followed by DNAse digestion (Promega) and purification using the RNeasy kit (Qiagen). Samples were sent for sequencing at the Earlham Institute (Norwich, UK) where 1ug of RNA was purified to extract mRNA with a poly-A pull down and 6 non-orientated libraries (LIB949-LIB954) constructed using the Illumina TruSeq RNA Library Preparation kit following manufacturer’s instructions. After cDNA synthesis 10 cycles of PCR were performed to amplify the fragments. Libraries were then pooled and sequenced on a single HiSeq 2000 lane generating 100bp paired-end sequences. Additionally to aid gene annotation a directional library (LIB1777) was constructed with RNA isolated from a mixture of asexual females at various developmental stages. Libraries were generated following the strand specific RNA sequencing method published by The Broad Institute [63], and sequenced to 100bp on a paired-end flow cell on the Illumina HiSeq2000 (Illumina, USA) (Additional File 21: Table S7).

### *Construction of a small RNA library of* M. persicae

RNA was extracted from 450 *M. persicae* nymphs using Tri-Reagent (Sigma). A small RNA library was prepared following the Illumina Small RNA v1.5 Sample Preparation protocol (Illumina Inc, San Diego, USA). Ligation of the 5` and 3` RNA adapters were conducted with 1μg RNA according to the manufacturers instructions (except that PCR was performed with 10mM dNTP in a 25μl reaction). Following ligation of the 5` and 3` RNA adapters, cDNA synthesis and PCR amplification, fragments corresponding to adapter-sRNA-adapter ligations (93-100bp) were excised from polyacrylamide gels and eluted using the manufacturer’s instructions. Sequencing was performed at The Sainsbury Laboratory (TSL, Norwich, UK) for 36nt single-end sequencing on an Illumina Genome Analyzer.

### Genome assembly and annotation

Full details of genome assembly, annotation and quality control are given in Additional File 2. Briefly, the genomes of *M. persicae* clones G006 and clone O were independently assembled using a combination of short insert paired-end and mate-pair libraries (Additional File 1: Table S1). Clone G006 was assembled with ALLPATHS-LG [64] and Clone O with ABySS [65] followed by scaffolding with SPPACE [66] and gapclosing with SOAP GapCloser [67]. Repetitive elements were annotated in both genomes with the REPET package (v2.0). We then predicted protein coding genes for each genome using the AUGUSTUS [68] and Maker [69] gene annotation pipelines using protein, cDNA and RNA-Seq alignments as evidence. A set of integrated gene models was derived from the AUGUSTUS and Maker gene predictions, along with the transcriptome and protein alignments, using EVidenceModeler [70]. Splice variants and UTR features were than added to the integrated EVidenceModeler predicted gene set using PASA [71]. Following these automatic gene annotation steps, manual annotation was performed for genes involved metabolism pathways and a subset of gene families implicated in host adjustment (Additional File 3).

### Gene family clustering

To investigate gene family evolution across arthropods we compiled a comprehensive set of proteomes for 17 insect lineages plus the branchiopod outgroup *D. pulex* and the spider mite *Tetranychus urticae* and combined them with the proteomes of the two newly sequenced *M. persicae* clones. In total 22 arthropod proteomes were included with all major insect lineages with publicly available genome sequences represented (Additional File 22: Table S8). In cases where proteomes contained multiple transcripts per gene the transcript with the longest CDS was selected. Although both *M. persicae* clones were included for clustering, comparisons between species were made using the G006 reference only. Putative gene families within our set of proteomes were identified based on Markov clustering of an all-against-all BLASTP search using the Markov Cluster Algorithm v.12.068 (MCL) [27]. Blast hits were retained for clustering if they had an E-value less than 1e^-5^ and if the pair of sequences aligned over at least 50% of the longest sequence in the pair. MCL was then run on the filtered blast hits with an inflation parameter of 2.1 and filtering scheme 6.

To estimate species phylogeny, protein sequences for 66 single copy conserved orthologs were extracted. For each gene, proteins were aligned using muscle v. 3.8.31 [72] followed by removal of poorly aligned regions with trimAl v. 1.2 [73]. The curated alignments were then concatenated into a supermatrix. Phylogenetic relationships were estimated using maximum likelihood (ML) in RAxML v. 8.0.23 [28]. The supermatrix alignment was partitioned by gene and RAxML was run with automatic amino acid substitution model selection and gamma distributed rate variation for each partition. One hundred rapid bootstrap replicates were carried out followed by a thorough ML tree search. As the focus of the present study is not on estimating absolute dates of divergence we used RelTime [29] to estimate the relative divergence times for species using the RAxML topology to generate an ultrametric phylogeny to use in the comparative analysis. RelTime has been shown to give relative dates of divergence that are well correlated with absolute divergence times derived from the most advanced Bayesian dating methods and is computationally tractable with large genomic datasets [29]. We estimated relative divergence times for species treating the supermatrix a single partition. RelTime was run with an LG model of protein evolution and the few clocks option (clocks merged on 2 std. errors).

### Analysis of gene family evolution

Gene family evolution across arthropods was investigated using CAFE v.3.0 [74]. CAFE models the evolution of gene family size across a species phylogeny under a maximum likelihood (ML) birth death model of gene gain and loss and simultaneously reconstructs maximum likelihood ancestral gene family sizes for all internal nodes, allowing the detection of expanded gene families within lineages. We ran CAFE on our matrix of gene family sizes generated by MCL under a birth death model of gene family evolution and modeled their evolution along the RelTime species tree. CAFE assumes that gene families are present in the last common ancestor of all species included in the analysis. To avoid biases in estimates of the rate of gene gain and loss we therefore removed gene families not inferred to be present in the last common ancestor of all taxa in the analysis based on maximum parsimony reconstruction of gene family presence / absence. Initial runs of CAFE produced infinite likelihood scores due to very large changes in family size for some gene families. We therefore excluded gene families where copy number varied between species by more than 200 genes. In total 4,983 conserved gene families were included for analysis. To investigate variation in the rate of gene birth and death (λ) across the arthropod phylogeny we tested a series of nested, increasingly complex, models of gene family evolution using likelihood ratio tests [75]. Models tested ranged from one with a single λ parameter across the whole phylogeny to a model with separate λ parameters for each of the major arthropod groups and a separate rate for each aphid species (Additional File 23: Table S9). For a more complex model to be considered an improvement a significant increase in likelihood had to be observed (likelihood ratio test, p < 0.05). For the best fitting model of gene family evolution (‘clade specific rates’, Additional File 24: Table S10), the average per gene family expansion and the number of expanded families were compared for each taxon included in the analysis. To correct for evolutionary divergence between taxa, average per gene family expansion and the number of expanded gene families were normalised for each taxon by dividing by the relative divergence time from the MRCA of the taxon in question (RelTime tree, branch length from tip to first node).

### Aphid gene duplication history and patterns of molecular evolution

To investigate the history of gene duplication in aphids we reconstructed the complete set of duplicated genes (paralogs) in *M. persicae* and *A. pisum* and calculated the rates of synonymous substitution per synonymous site (*d_S_*) and non-synonymous substitution per non-synonymous site (*d*_*N*_) between each duplicated gene and its most recent paralog. We then created age distributions for duplicate genes in the two aphid genomes based on *d_S_* values between paralogs and compared rates of evolution based on *d*_*N*_/_*_dS_*_ ratios. Larger values of *d_S_* represent older duplication events, and the *d*_*N*_/*d_S_* ratio reflects the strength and type of selection acting on the sequences. Paralog pairs were identified by conducting an all-against-all protein similarity search with BLASTP on the proteome of each species with an E-value cutoff of e^−10^. When multiple transcripts of a gene were present in the proteome the sequence with the longest CDS was used. Paralogous gene pairs were retained if they aligned over at least 150 amino acids with a minimum of 30% identity [76]. For each protein only the nearest paralog was retained (highest scoring BLASTP hit, excluding self hits) and reciprocal hits were removed to create a non-redundant set of paralog pairs. For each paralog pair a protein alignment was generated with muscle v. 3.8.31 [72]. These alignments were then used to guide codon alignments of the CDS of each paralog pair using PAL2NAL [77]. From these codon alignments pairwise *d*_*N*_ and *d_S_* values were calculated with paml v4.4 using YN00 [78]. Paralog pairs with *d_S_* > 2 were excluded from our analysis as they likely suffer from saturation. For the generation of age distributions we used all gene pairs that passed our alignment criteria. For comparisons of rates of evolution (*d*_*N*_/*d_S_*) we applied strict filtering criteria to avoid inaccurate *d*_*N*_/*d*_*S*_ estimates caused by insufficiently diverged sequences; pairs were removed if they had *d*_*N*_ or *d_S_* less than 0.01 and fewer than 50 synonymous sites. We also calculated pairwise *d*_*N*_ and *d_S_* for 1:1 orthologs between *M. persicae* and *A. pisum* (extracted from the MCL gene families). This allowed us to separate duplicated genes into ‘old’ (before speciation) and ‘young’ (after speciation) categories depending on whether *d_S_* between a paralog pair was larger or smaller than the mean *d_S_* between 1:1 orthologs which corresponds to the time of speciation between the two aphid species. Adding 1:1 orthologs also allowed us to compare rates of evolution (*d*_*N*_/*d_S_*) between single copy and duplicated genes. In addition to the pipeline above, we also identified tandemly duplicated genes in the *M. persicae* genome using MCSscanX [79].

### *RNA-seq analysis of* M. persicae *clone O colonies on different plant species*

To identify genes involved in *M. persicae* host adjustment we compared the transcriptomes of clone O colonies reared on either *B. rapa* or *N. benthamiana* for one year (LIB949 – LIB954, Additional File 21: Table S7). Reads were quality filtered using sickle v1.2 [80] with reads trimmed if their quality fell to below 20 and removed if their length fell to less than 60 bp. The remaining reads were mapped to the G006 reference genome with Bowtie v1.0 [81] and per gene expression levels estimated probabilistically with RSEM v1.2.8 [82]. We identified differentially expressed genes with DEseq [83] using per gene expected counts for each sample generated by RSEM. To increase statistical power to detect differentially expressed genes, lowly expressed genes falling into the lowest 40% quantile were removed from the analysis. Genes were considered differentially expressed between the two treatments if they had a significant p value after accounting for a 10% false discovery rate according to the Benjamini-Hochberg procedure and if a fold change in expression of at least a 1.5 was observed.

### qRT-PCR analyses

Total RNA was isolated from adults using Trizol reagent (Invitrogen) and subsequent DNase treatment using an RNase-free DNase I (Fermentas). cDNA was synthesised from 1 μg total RNA with RevertAid First Strand cDNA Synthesis Kit (Fermentas). The qRT-PCRs reactions were performed on CFX96 Touch™ Real-Time PCR Detection System using gene-specific primers (Additional File 25: Table S11). Each reaction was performed in a 20 μL reaction volume containing 10μL SYBR Green (Fermentas), 0.4 μL Rox Reference Dye II, 1 μL of each primers (10 mM), 1 μL of sample cDNA, and 7.6 μL UltraPure Distilled water (Invitrogen). The cycle programs were: 95°C for 10 s, 40 cycles at 95°C for 20 s, 60°C for 30 s. Relative quantification was calculated using the comparative 2^−ΔCt^ method [84]. All data were normalised to the level of *Tubulin* from the same sample. Design of gene-specific primers were achieved by two steps. First, we used PrimerQuest Tool (Integrated DNA technologies, Iowa USA) to generate five to ten qPCR primer pairs for each gene. Then, primer pairs were aligned against cathepsin B and cuticular protein genes. Only primers aligned to unique sequences were used (Additional File 25: Table S11). Genes for which no unique primers could be designed were excluded from analyses.

### Plant host switch experiments

The *M. persicae* clone O colony reared on *B. rapa* was reared from a single female and then transferred to *A. thaliana* and *N. benthamiana* and reared on these plants for at least 20 generations. Then, 3^rd^ instar nymphs were transferred from *A. thaliana* to *N. benthamiana* and vice versa for three days upon which the insects were harvested for RNA extractions and qRT-PCR analyses.

### Cloning of dsRNA constructs and generation of transgenic plants

A fragment corresponding to the coding sequence of MpCathB4 (Additional File 18) was amplified from *M. persicae* cDNA by PCR with specific primers containing additional attb1 (ACAAGTTTGTACAAAAAAGCAGGCT) and attb2 linkers (ACCACTTTGTACAAGAAAGCTGGGT) (MpCathB4 attB1 and MpCathB7 attB2, Additional File 25: Table S11) for cloning with the Gateway system (Invitrogen). A 242-bp MpCathB4 fragment was introduced into pDONR207 (Invitrogen) plasmid using Gateway BP reaction and transformed into DH5α. Subsequent clones were sequenced to verify correct size and sequence of inserts. Subsequently, the inserts were introduced into the pJawohl8-RNAi binary silencing vector (kindly provided by I.E. Somssich, Max Planck Institute for Plant Breeding Research, Germany) using Gateway LB reaction generating plasmids pJMpCathB4, which was introduced into *A. tumefaciens* strain GV3101 containing the pMP90RK helper plasmid for subsequent transformation of *A. thaliana* using the floral dip method [85]. Seeds obtained from the dipped plants were sown and seedlings were sprayed with phosphinothricin (BASTA) to selection of transformants. F2 seeds were germinated on Murashige and Skoog (MS) medium supplemented with 20 mg ml BASTA for selection. F2 plants with 3:1 dead/alive segregation of seedlings (evidence of single insertion) were taken forward to the F3 stage. Seeds from F3 plants were sown on MS+BASTA and lines with 100% survival ratio (homozygous) were selected. The presence of pJMpCathB4 transgenes was confirmed by PCR and sequencing. Three independent pJMpCathB4 transgenic lines were taken forward for experiments with aphids. These were At_dsCathB 5-1, 17-5 and 18-2.

To assess if the 242-bp MpCathB4 fragment targets sequences beyond cathepsin B genes, 242-bp sequence was blastn-searched against the *M. persicae* clones G006 and O predicted transcripts at AphidBase and cut-off e-value of 0.01. The sequence aligned to nucleotide sequences of MpCathB1 to B13 and MpCathB17 with the best aligned for MpCathB4 (241/242, 99% identity) and lowest score for MpCathB17 (74/106, 69% identity) (Additional File 18). *M. persicae* fed on At_dsCathB 5-1, 17-5 and 18-2 transgenic lines had lower transcript levels of AtCathB1 to B11, whereas that of MpCathB12 was not reduced (Fig. 4.1A). Identity percentages of the 242-bp fragment to AtCathB1 to B11 range from 99% to 77%, whereas that of MpCathB12 is 73% (Additional File 18). Thus, identity scores higher than 73-77% are needed to obtain effective RNAi-mediated transcript reduction in *M. persicae*.

### Plant-mediated RNA interference (RNAi) of GPA cathepsin B genes

Seed of the pJMpCathB4 homozygous lines (expressing dsRNA corresponding to Cathepsin B, dsCathB, Additional File 18) was sown and seedlings were transferred to single pots (10 cm diameter) and transferred to an environmental growth room at temperature 18°C day/16°C night under 8 hours of light. The aphids were reared for 4 generations on *A. thaliana* transgenic plants producing dsGFP (controls) and dsCathB. Five *M. persicae* adults were confined to single 4-week-old *A. thaliana* lines in sealed experimental cages (15.5 cm diameter and 15.5 cm height) containing the entire plant. Two days later adults were removed and five nymphs remained on the plants. The number of offspring produced on the 10th, 14th, 16th day of the experiment were counted and removed. This experiment was repeated three times to create data from three independent biological replicates with four plants per line per replicate.

## Data availability

All assemblies and annotation features are available and downloadable at www.aphidbase.com [86]. Sequence data has been deposited in the sequence read archive at the European Nucleotide Archive (ENA) and are available under BioProject accessions PRJEB11304 (clone O) and PRJNA319804 (G006).

## Acknowledgements

We thank Brain Fenton for help with genotyping *M. persicae* clone O, Linda M. Field for being a co-investigator on the Capacity and Capability Challenge (CCC-15) project that funded the first round of genome sequencing of *M. persicae* clone O. We are grateful to Ian Bedford and Gavin Hatt (JIC Insectary) for rearing and care of aphids and the John Innes Horticultural Services for growing the plants used in this study. Next-generation sequencing and library construction was delivered via the BBSRC National Capability in Genomics (BB/J010375/1) at the Earlham Institute (formerly The Genome Analysis Centre), Norwich, by members of the Platforms and Pipelines Group.

## Funding

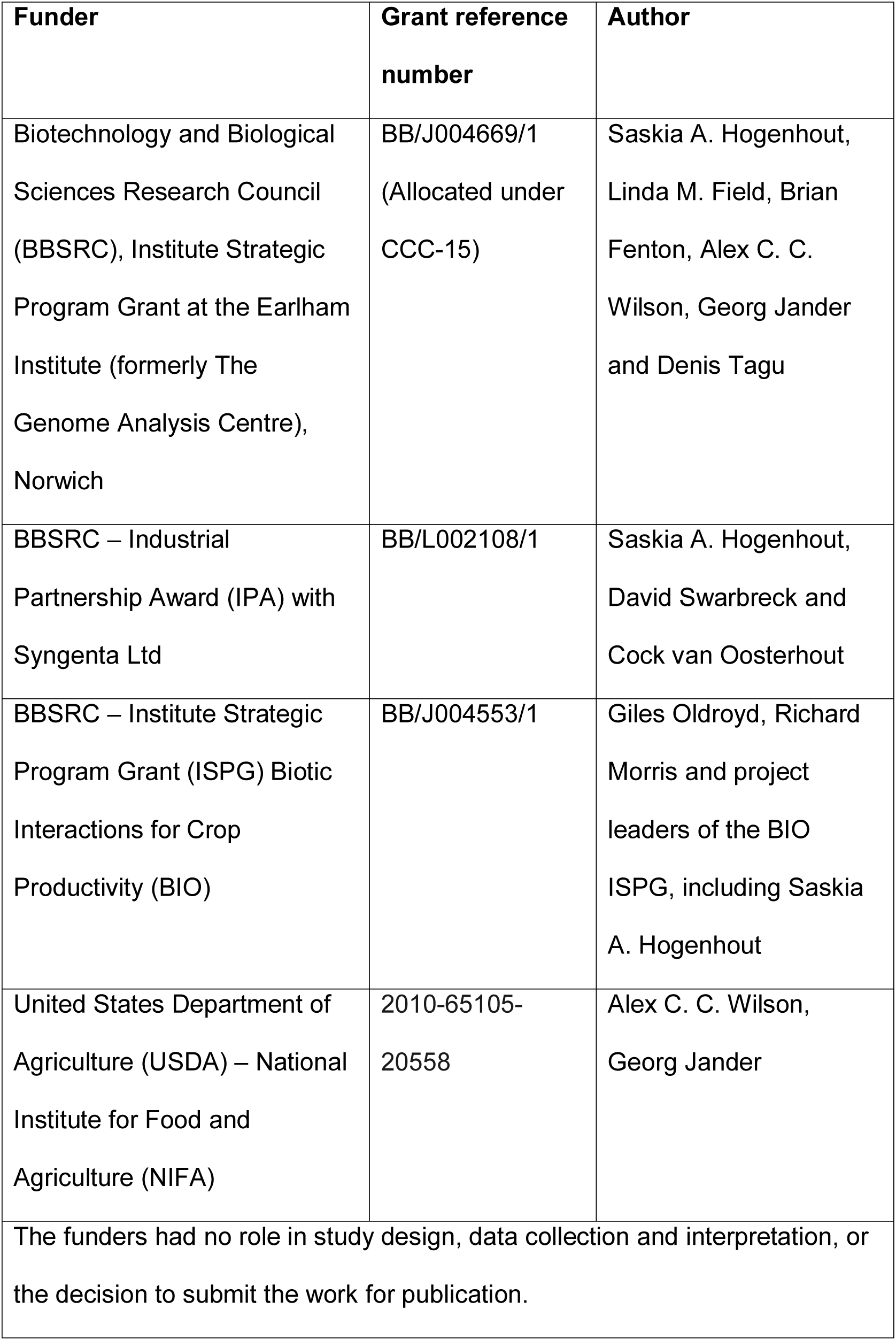

## Additional Files

### Additional File 1: Table S1

Format: .docx

Summary of libraries generated and datasets used for assembly of the genomes of *Myzus persicae* clones G006 and O.

### Additional File 2

Format: .docx

Supplementary Text: Genome assembly, annotation and quality control. 1230

### Additional File 3

Format: .docx

Supplementary Text: Annotation of metabolic processes and specific gene families. 1234

### Additional File 4

Format: .docx

Figure S1: Maximum likelihood phylogeny of 21 arthropod species with fully sequenced genomes based on 66 strictly conserved singly copy orthologs. Sequences were aligned with MUSCLE [72] and trimmed to remove poorly aligned regions with TrimAl [73]. The phylogeny was estimated using RAxML [28] with each gene treated as a separate partition. Automatic protein model selection was implemented in RAxML with gamma distributed rate variation. Values at nodes show bootstrap support based on 100 rapid bootstrap replicates carried out with RAxML.

### Additional File 5

Format: .docx

Figure S2: Enriched GO terms relating to biological processes of aphid specific *M. persicae* genes. GO term enrichment analysis was carried out using Fishers’ exact test in BINGO [89] with correction for multiple testing applied by the Benjamini-Hochberg procedure allowing for a 10% false discovery rate. GO terms from aphid specific genes were compared to GO terms from the complete set of *M. persicae* genes. Enriched GO terms were reduced and visualised with REVIGO [90]. GO terms are clustered by semantic similarity with the size of each circle relative to the size of the GO term in UniProt (larger circles = more general GO terms) and coloured by their p values according to the Fisher exact test of enrichment. A complete list of enriched GO terms for aphid specific genes are given in Additional File 6: Table S2.

### Additional File 6

Format: .xlsx

Table S2: Overrepresented GO terms in aphid specific genes compared to the genome as a whole. Overrepresented GO terms identified with BINGO [89] using Fisher’s exact test accounting for a 10% false discovery rate.

### Additional File 7

Format: .docx

Figure S3: Model based analysis of gene gain and loss across arthropods. Gene gain and loss (λ) was modelled across the arthropod phylogeny under a birth-death process with CAFE [74] for 4,983 widespread gene families inferred to be present in the most recent common ancestor (MRCA) of all included taxa. Nested models with increasing numbers of lambda parameters were compared using likelihood ratio test (Additional File 23: Table S9; Additional File 24: Table S10). Results are shown for the best fitting clade specific rates model. **A)** Arthropod phylogeny scaled by clade specific ML values of λ inferred by CAFE. Branch colours indicate where separate λ parameters were specified. *A. pisum* (red) has undergone a significant increase in the rate of gene gain and loss (λ) compared to other arthropod species. **B)** Linear regression line (mean and 5-95% confidence intervals) of the number of expanded families versus the gain in gene number per family across arthropod taxa. Both the size of the expansion and the number of expanded families were log_10_ transformed and scaled relative to the divergence time. There is a significant positive relationship across arthropod taxa in the number of families that expand, and the mean number of genes gained within the expanded families (Regression: R^2^=29.4%; F_1,19_=9.32, p=0.007). The specialist aphid *A. pisum* (red) is an outlier, showing a relative excess in both the number of expanded families and the magnitude of the mean family expansion. In contrast, although the generalist aphid *M. persicae* (green) has many expanded families, it shows relatively little gene gain per family.

### Additional File 8

Format: .xlsx

Table S3: MCL gene families significantly expanded according to the binomial test in: **A)** both aphid species, **B)** *M. persicae* but not *A. pisum*,**C)** *A. pisum* but not *M. persicae*. The number of genes per MCL gene family for each species included for gene family clustering is given. p values were calculated for each of the two aphid species, for each MCL family, by comparing the number of members in *M. persicae* or *A. pisum* to a binomial distribution drawn from the mean family size excluding aphids. A p value less than 0.05 was considered to imply a significant gene family expansion. In total 6,148 MCL families found in both aphid species and at least one other taxon were tested. MCL gene families are ordered by prevalence in taxa other than aphids with the most widespread families listed first. MCL families with at least one *M. persicae* member differentially expressed in the host swap RNA-seq experiment are highlighted in yellow. Each expanded MCL family is annotated with expression information for *M. persicae* family members in the host swap RNA-seq experiment (averaged across all 6 RNA-seq libraries and expressed in fragments per kilobase of transcript per million mapped reads (FPKM)), number and proportion of *M. persicae* members differentially expressed in the host swap RNA-seq experiment, InterProScan domains and descriptions, InterProScan GO terms and Blast2GO GO terms and descriptions. All families are annotated based on *M. persicae* members.

### Additional File 9

Format: .docx

Figure S4: The rate of evolution (*d_N_*/*d_S_*) versus time since duplication (*d_S_*) for *A. pisum* and *M. persicae* paralog pairs. Paralog pairs that duplicated before the divergence of *A. pisum* and *M. persicae* (*d_S_* > 0.26) are coloured blue, paralog pairs that duplicated after the divergence of *A. pisum* and *M. persicae* (*d_S_* < 0.26) are coloured red.

### Additional File 10

Format: .docx

Supplementary Text: *M. persicae* phylome report. 1320

### Additional File 11

Format: .docx

Table S4: *M. persicae* genes that are differentially expressed in aphids reared on different host plants. Genes that show differential expression (>1.5-fold with 10% FDR) on *Nicotiana benthamiana* vs. *Brassica rapa* (Nb/Br) are listed. Top: genes more highly expressed on *B. rapa,* and bottom: more highly expressed on *N. benthamiana.* Fold-change is the average over 3 biological replicates on each host plant, *p*-value and FDR-adjusted-*p*-value (padj) based on DE-seq analysis. Annotations were conducted by NCBI blastX using cDNA sequences. FC is fold-change of Nb expression vs. Br. The presence of a predicted secretory signal peptide is indicated with “*” in the SP column. Tissue-specific expression is indicated with “+” in the “SG” (salivary gland), “Gut” and “Head” columns, based on detection of the sequence in tissue-specific EST data [23].

### Additional File 12

Format: .docx

Figure S5: The Rebers and Riddiford subgroup 2 (RR-2) cuticular protein genes that are differentially expressed upon *M. persicae* host change belong predominantly to a single aphid-expanded clade and form gene clusters in the *M. persicae* genome. **A)** Maximum likelihood phylogenic tree of arthropod RR-2 cuticular protein protein sequences. The sequences were aligned with Muscle [72] and the phylogeny estimated using FastTree [88] (JTT + CAT rate variation). Circles on branches indicate SH-like local support values >80%, scale bar below indicates 0.1 substitutions per site. Rings from outside to inside: ring 1, *M. persicae* RR-2 cuticular protein (MpCutP) gene identities (IDs) with numbers in red indicating upregulation of these genes in *M. persicae* reared for one year on *N. benthamiana* relative to those reared for one year on *B. rapa*, and bold font indicating location on the RR-2 cuticular protein multigene clusters shown in **B**; ring 2, red squares indicating MpCutP genes that are differentially expressed upon *M. persicae* host change; ring 3, CutP genes from different arthropods following the color scheme of the legend in the lower left corner and matching the colors of the branches of the phylogenetic tree; ring 4, aphid expanded (AE) clades with AE_Clade I labeled light green and AE_Clade II light blue. **B)** MpCutP multigene clusters of the *M. persicae* genome. Lines indicate the genomic scaffolds on which the MpCutP genes are indicated with block arrows. Gene IDs above the genes match those of the phylogenetic tree in A, with block arrows and fonts highlighted in red being DE upon host change. Scale bar on right shows 20 kb.**C)** Relative expression levels of MpCutP genes of *M. persicae* at 7 weeks being reared on *N. benthamiana* (Nb), *B. rapa* (Br) and *A. thaliana* (At). Numbers under the graphs indicate MpCutP gene IDs with those in red font differentially expressed as in A. Batches of five adult females were harvested for RNA extraction and qRT-PCR assays. Bars represent expression values (mean ± standard deviation (SD)) of three independent biological replicates. \**p*<0.05 (Anova with Fishers LSD to control for multiple tests). **D)** As in C, except that individual aphids reared on At were transferred to At (At to At) or Nb (At to Nb) and harvested at 2 days upon transfer. **E**) As in D, except that individual aphids reared on Nb were transferred to Nb (Nb to Nb) or At (Nb to At) and harvested at 2 days upon transfer.

### Additional File 13

Format: .docx

Figure S6: UDP-glucosyltransferase (UGT) genes that show differential expression on different host plants fall within an aphid specific clade, and some are associated with an array of tandem duplicates. Phylogeny of (UGT) proteins from *M. persicae* (green), *A. pisum* (blue), *R. prolixus* (purple) and *D. melanogaster* (red). Protein sequences were aligned with Muscle [72] and the phylogeny estimated using RAxML [28] with automatic model selection and gamma distributed rate variation. 100 rapid bootstrap replicates were carried out with RAxML. Grey circles on branches indicate bootstrap support greater than 80%. Genes showing elevated expression in aphid reared on *B. rapa* are indicated in red. Bottom, part of scaffold_555 containing 7 predicted UGT genes, 4 of which are more highly expressed on *B. rapa* host plants. Scale bar at left is 10 kb.

### Additional File 14

Format: .docx

Figure S7: Maximum likelihood phylogeny of Cytochrome-P450 proteins from *M. persicae* (green), and *A. pisum* (blue). *A. pisum* P450s are named according to their annotation from the Cytochrome-P450 homepage [91]. Protein sequences were aligned with Muscle [72] and the phylogeny estimated using RAxML [28] with automatic model selection and gamma distributed rate variation. 100 rapid bootstrap replicates were carried out with RAxML. Grey circles on branches indicate bootstrap support greater than 80%. Transcripts that show elevated expression on *B. rapa* are indicated with red arrowhead, and on *N. benthamiana* with a green arrowhead. Bottom: scaffold 338 containing 4 differentially expressed cytochrome-P450 genes (red), together with 3 non-regulated p450s (white) is shown. (*locus 000111270 is one of 8 *M. persicae* P450 genes that were excluded from phylogenetic analysis after manual curation as they were either fragmented or had incorrect annotations.

### Additional File 15

Format: .docx

Figure S8: Maximum likelihood phylogeny of *M. persicae*, *A. pisum* and *D. melanogaster* lipase-like genes (MCL family 16). Protein sequences were aligned with Muscle [72] and the phylogeny estimated using RAxML [28] with automatic model selection and gamma distributed rate variation. 500 rapid bootstrap replicates were carried out with RAxML, bootstrap support values are shown at nodes. Genes showing significantly elevated expression on *B. rapa* are indicated with red arrows. Below, part of scaffold 351 which contains 4 lipase-like genes in a tandem array, 3 of which are up-regulated in aphids reared on *B. rapa*.

### Additional File 16

Format: .xlsx

Table S5: Rates of evolution for all *M. persicae* paralog pairs containing at least one DE gene between *M. persicae* clone O reared for 1 year on *B. rapa* or *N. benthamiana*. Pairwise *d_N_*/*d_S_* was estimated using the YN00 [87] model in PAML [78]. Paralog pairs or ordered by *d_S_* (youngest duplicates first). Light green shading indicates duplication after speciation of *M. persicae* and *A. pisum* (*d_S_* < 0.26). Paralog pairs coloured red have 0 *d_S_* and *d_N_* (identical coding sequences).

### Additional File 17

Format: .docx

Figure S8.5: Performance of *M. persicae* clone O on three plant species. Three one-day old *M. persicae* clone O nymphs were placed on the youngest leaf of each of the following plant species: *Arabidopsis thaliana;* (At); *Brassica rapa* (Br); and *Nicotiana benthamiana* (Nb)]. The aphids were then examined for developmental time, ^1423 reproduction rate, longevity, and weight. The developmental time is^ the duration in days between birth of the nymph and emergence of the adult. The reproduction rate is the number of progeny produced by a single adult female over a period of 10 days. The weight is measured for 10 adult females on the first day they become adults. The longevity is the number of days between birth and death. Columns represent values (mean ± SD) from 10 independent biological replicates (*p* >0.05).

### Additional File 18

Format: .docx

Cathepsin B dsRNA alignment. Blast search results of the cathepsin B dsRNA sequence used to generate transgenic lines for plant-mediated RNAi of GPA. The 242-bp fragment of MpCathB4 (Clone O) was blastn-searched against the annotated genome of *M. persicae* clones G006. Identities and gaps of 242 bp with MpCathB4, MpCathB5, MpCath10 and MPCathB11 (indicated with *) were generated by NCBI blastn suite-2 sequences because these were misannotated in MyzsDB (G006).

### Additional File 19

Format: .docx

Figure S9: Cathepsin B expression levels of CathB-RNAi and control (dsGFP-exposed) aphids after two days on non-transgenic *A. thaliana* and *N. benthamiana* plants. Ten 3^rd^ instar nymphs on *dsCathB* (lines 17-5 and 18-2) and *dsGFP* transgenic *A. thaliana* lines were transferred to non-transgenic *A. thaliana* (At) (**A**) and non-transgenic *N. benthamiana* (Nb) (**B**) plants. Aphids were harvested two days later for RNA extraction and qRT-PCR analyses. Bars represent mean ± SD of the relative *M. persicae* CathB expression levels (compared to aphids on *dsGFP* (control) plants) of three independent biological replicates with five adult females each. \**p*<0.05.

### Additional File 20

Format: .docx

Figure S10: Domain analysis of aphid-specific cathepsin B genes. Protein sequences were used for analysis in InterPro. Clade highlighted in light green aphid-specific clade I, and the blue is aphid-specific clade II. Asterisks (*) indicate cathepsin B with complete domains, green asterisks are *M. persicae* cathepsins B, blue asterisks are the *A. pisum* ones, and a red asterisk are the *D. noxia* ones. 1458

### Additional File 21

Format: .docx

Table S7: Summary of all newly generated *M. persicae* RNA-seq transcriptome data. SS = Strand-specific RNA-seq reads

### Additional File 22

Format: .xlsx

Table S8: Proteomes of 22 arthropod genomes used for comparative gene family analyses. 1468

### Additional File 23

Format: .docx

Table S9: Models tested in the CAFE analysis of gene family evolution. Increasingly complex models of gene family evolution were tested using CAFE [74] with a focus on determining if aphid rates of gene gain and loss (gain=loss=λ) differ from that of other arthropod lineages. Regions of the arthropod phylogeny with differentλ parameters were specified with the λ tree (newick format), which follows the species tree. For each model, 5 runs were conducted to check convergence. F.P. = free parameters, Lh. = likelihood, S.D. = standard deviation.

### Additional File 24

Format: .docx

Table S10: Likelihood ratio test results comparing models of gene family evolution estimated in CAFE [74]. Models tested are detailed in Additional File 23: Table S9. Likelihood ratio tests were conducted in a nested fashion comparing more complex models to less complex models. The best fitting model tested was the clade specific rates model, which gave a significant increase in likelihood over all other more simple models. The difference in the number of free parameters between each model is shown below the shaded squares. The likelihood ratio and p value (in brackets) for each model comparison are shown to the right of the shaded squares. For each model the best likelihood score out of five runs was used to calculate the likelihood ratio. The likelihood ratio was calculated as follows: likelihood ratio = 2 × ((likelihood more complex model) − (likelihood less complex model)). p values for the likelihood ratio test were generated by comparing the likelihood ratio between the more complex model and the less complex model to a chi square distribution with the degrees of freedom equal to the difference in the number of free parameters between the two models.

### Additional File 25

Format: .docx

Table S11: Sequences of primers used in Gateway cloning and qRT-PCR experiments.

### Additional File 26

Format: .docx

Table S12: List of cathepsin B genes annotated in the genome of the pea aphid *Acyrthosiphon pisum.*

### Additional File 27

Format: .docx

Table S13: List of cathepsin B genes annotated in the genomes of *Myzus persicae* clones G006 and 0. Four fragments were annotated as MpCathB1 and two as MpCathB3 in Clone O. Alignment of fragments to corresponding genes indicated that fragments were part of the gene. The sequences of MpCathB1 and MpCathB3 in clone O were confirmed by PCR and sequencing. MpCathB2, MpCathB7, and MpCathB12 were missing in the genome annotation of *M. persicae* clone O; their sequences were confirmed by PCR and sequencing.

### Additional File 28

Format: .xlsx

Table S14: Cathepsin B genes for 26 arthropod species. Cathepsin B sequences were previously annotated for *A. pisum* by Rispe et al. (53). These sequences all fall into MCL family_110. Additional Cathepsin B sequences were identified for *N. lugens*, *D. citri*, *D. noxia* and *M. destructor* based on blastp similarity searches to *M. persicae* clone G006 cathepsin B sequences (see sub tables A-D). Sequences coloured red were considered fragments and excluded from subsequent phylogenetic analysis.

### Additional File 29

Format: .xlsx

Table S15: Annotation of cuticular proteins in 5 hemipteran species. A) Overview of cuticular protein family size in 5 hemipteran genomes. Cuticular proteins were identified using CutProtFamPred on the proteomes of *M. persicae* clone G006, *A. pisum*, *D. noxia*, *D. citri*, *N. lugens* and *R. prolixus*. The number of genes DE between *M. persicae* clone O individuals reared on either *B. rapa* or *N. benthamiana* is also given for each family. Full results of the CutProfFamPred analysis for all 5 proteomes are given in B). The RR2 cuticular protein family thas the highest number of genes DE between aphids reared on *B. rapa* and *N. benthamiana* and was subjected to phylogenetic analysis. Due to the high variability of RR2 cuticular proteins phylogenetic analysis was carried out on the RR2 domain only. 8 genes were removed from the analysis due to poor alignment of the RR2 domain. C) blastp identification of the RR2 domain in RR2 cuticular proteins.

### Additional File 30

Format: .docx

Table S16: Summary of the manual annotation and gene edition of *M. persicae* (clone G006) CPR as described in Additional File 3.

### Additional File 31

Format: .xlsx

Table S17: P450 genes identified in *M. persicae* clone G006. P450 genes were identified based on blastp searches against *A. pisum* P450 sequences obtained from the P450 website [91] and presence of the PF00067 P450 domain. Genes were manually checked for completeness and missanotated genes removed from the phylogenetic analysis (highlighted in red).

### Additional File 32

Format: .xlsx

Table S18: Functional annotation of *M. persicae* clone G006 lipase-like proteins found in MCL family_16.

### Additional File 33

**Table S19:** UDP-glucosyltransferase (UGT) genes found in *M. persicae* clone G006, *A. pisum*, *R. prolixus* and *D. melanogaster*. UGT genes were identified by searching the MCL gene families for known *D. melanogaster* UGT proteins listed in FlyBase (www.flybase.org/). All annotated *D. melanogaster* UGT genes were found in a single MCL genes family (family_12).

